# Severe feed restriction in a fish: is the positive effects of nucleotide supplementation on growth recovery explained by one-carbon metabolism?

**DOI:** 10.64898/2026.06.03.729921

**Authors:** Charles Drouin-Johnson, Nathalie R. Le François, Grant W. Vandenberg, Moïse Cantin, Simon G. Lamarre

**Affiliations:** Laboratoire de physiologie et aquaculture de la conservation, Collections vivantes, recherche et développement scientifique, Biodôme de Montréal/Espace pour la vie, 4777, Ave Pierre-De Coubertin, Montréal, QC Canada H1V 1B3; Département des sciences animales, Pavillon Paul-Comtois, 2425, rue de l’Agriculture, Local 4131, Université Laval, Québec, QC Canada G1V A06; Pisciculture-des-Monts-de-Bellechasse Inc., 251, rue Commerciale, Saint-Damien-de-Buckland, QC, Canada G0R 2Y0; Département de biologie, Université de Moncton, 18, ave. Antonine-Maillet, Moncton, NB, Canada E1A 3E9

**Keywords:** Fasting, Compensatory growth, 1C metabolism, dietary nucleotide, *Salvelinus fontinalis* × *Salvelinus alpinus*

## Abstract

Charrs (*Salvelinus* spp.) are strong candidates for cyclical feed-restriction strategies due to their documented exceptional ability to withstand and recover from feed resources scarcity during winter (Drouin-Johnson et al. 2026; Le François et al. 2023; Savoie et al. 2017). Dietary nucleotides (NT) are supplements used to improve growth and health in fish, yet their integration into feed-restriction programs to enhance growth recovery remains largely unexplored. The one-carbon (1C) metabolism offers a mechanistic framework to investigate these effects, as it relies on vitamin B status and supports methylation reactions central to biosynthesis and growth. A large-scale trial involving *S. alpinus* × *S. fontinalis* was conducted, comprising a 90-day fasting period followed by refeeding on a commercial feed with or without NT. Growth and feeding metrics were evaluated alongside key metabolites of 1C metabolism to assess adaptations to fasting-induced vitamin B deficiency and the role of NT in recovery. Starved fish lost 5.2 % of body mass (*p* < 0.05) but displayed accelerated growth upon refeeding, further enhanced by NT supplementation by 27.8 % (*p* < 0.05), forecasting full compensation after 125 days. Starvation increased homocysteine and decreased methionine concentrations (*p* < 0.05), while S-adenosylmethionine, S-adenosylhomocysteine, and formate showed only minor fluctuations. These patterns indicate moderate but reversible fluctuations of 1C metabolism. NT supplementation may exert a methyl-unit sparing effect during refeeding, potentially supporting anabolic recovery. This study uniquely combines commercial-scale growth trials and molecular analyses, providing new insights into a species-adapted feeding strategy that enhances growth recovery performances and metabolic resilience.

## 1. Introduction

The Arctic charr (*Salvelinus alpinus*) is distributed across the Holarctic realm and beyond while the brook charr, *S. fontinalis,* is occurring from the Laurentian Plateau of northern Québec (Canada) to the southern Appalachians (USA). Both species co-occur in the northern range of brook charr and can experience extreme seasonal changes in water temperature, photoperiod, and food availability [1, 2]. These conditions have shaped remarkable adaptations, including the ability to endure prolonged food items scarcity during winter and to rapidly restore energy stores during short summer feeding periods [3, 4, 5]. Fasting imposes major metabolic shifts and activates a chronic stress response, characterized by three well-defined phases [6]; an initial transition (phase I), followed by lipid mobilization (phase II), and eventually protein catabolism (phase III), the latter associated with elevated stress and emaciation. Lipid-rich species such as charrs delay transition into phase III, making them especially resilient to feed deprivation. Arctic charr, the northernmost salmonid, may be among the most fasting-adapted vertebrates: mature anadromous females have been observed to forego feeding and reproduction over nearly year-long riverine periods [7, 8]. Adaptation to fasting in Arctic charr for example, has been investigated from multiple angles, including protein and glucose metabolism [3, 9], adiposity and appetite regulation [10, 11], and enzymatic activity [12, 13]. In captivity, the Arctic charr exhibits seasonal ‘voluntary anorexia,’ ceasing growth even under constant conditions [12, 13, 14, 15]. Studies by Striberny et al. [16, 17, 18] have shown that this trait is under tight endogenous control, distinguishing charrs from other salmonids and positioning *S. alpinus* as a valuable model to study nutrition-mediated energy homeostasis.

In aquaculture, optimizing growth by manipulating diet composition and feeding regime is a key strategy [19, 20], as feed costs can account for 50-70% of production expenses [21]. Recently, feed restriction trials have been piloted and two emerging strategies in Arctic charr farming are: (1) exploiting fat reserves to reduce feed and labor costs without long-lasting effects on productivity, and (2) inducing a robust compensatory growth (CG) response after a severe winter fast [22, 23]. CG, a well-documented phenomenon across taxa, enables rapid catch-up growth and offers promising applications for fish production [24]. In the present study the intergeneric hybrid *Salvelinus fontinalis* × *Salvelinus alpinus* was used based on its interest in aquaculture. Their geographical distributions overlap and hybridization occurs in the wild [25 26]. They display distinct genomic organization, chromosome rearrangement and phenotypes [27] and hybridization was reported to increase resistance to skin infections parasites outbreaks [28), and productivity traits in comparison to the parental species *S. fontinalis* [29].

In parallel, dietary nucleotide (NT) supplementation has been proposed to enhance post-fasting growth, possibly by acting as a chemoattractant [30], improving gut morphology [31], or sparing energy that would otherwise be used for *de novo* NT synthesis [32]. Although NT supplementation has shown growth benefits in several species, including Arctic charr [33], the underlying mechanisms remain unclear [34, 35].

One-carbon (1C) metabolism is ubiquitous in living cells and supports the transfer of methyl groups, which are essential for biosynthetic and regulatory processes, including *de novo* nucleotide synthesis, amino acid metabolism, and DNA and protein methylation [36, 37]. In animals, methyl units must be obtained through the diet, and nutritional substrates and cofactors of 1C metabolism include folate, vitamins B_2_, B_6_, and B_12_, methionine (Met), choline, and betaine [38, 39]. These micronutrients are particularly relevant in salmonid aquaculture, where growth modulation and efficient metabolic adaptation are key production traits and where vitamin or methyl-donor deficiencies can directly impair protein synthesis, lipid metabolism, and immune function. Within this pathway, remethylation regenerates Met from homocysteine (Hcy), maintaining the supply of Met for S-adenosylmethionine (SAM) synthesis, the universal methyl donor for cellular methylation reactions [40, 41]. SAM is converted to S-adenosylhomocysteine (SAH) after donating its methyl group, and then to Hcy, which can be remethylated to Met or diverted through the transsulfuration pathway to form cysteine. Because SAM and SAH represent the substrate and product of methylation reactions, their ratio serves as a robust proxy for the methylation capacity and functional state of 1C metabolism [37, 42, 43].

As shown in Figure 1, one-carbon (1C) metabolism is compartmentalized across the cytosol, mitochondria, and nucleus [40]. Mitochondrial formate production provides the transferable 1C units that fuel cytosolic biosynthetic reactions, including de novo nucleotide synthesis [37, 44]. Because formate reflects mitochondrial 1C flux, plasma formate concentration has been proposed as a sensitive indicator of vitamin B-related impairments of 1C metabolism in mammals [42]. Such deficiencies have been linked to altered nervous and cellular function, reduced metabolic efficiency, and growth impairment in several animal models [44–46], yet have never been examined in fish. Despite charr’s remarkable capacity to withstand long periods of starvation, its ability to maintain methyl-unit balance and methylation potential during fasting remains unknown. Understanding these adjustments may reveal how this species sustains basal biosynthetic functions under severe nutrient limitation. Because 1C metabolism also supports anabolic recovery when feeding resumes, we hypothesize that dietary nucleotide (NT) supplementation could reduce the need for endogenous NT synthesis, thereby sparing methyl donors for other growth-related processes during refeeding.

**Figure 1.**
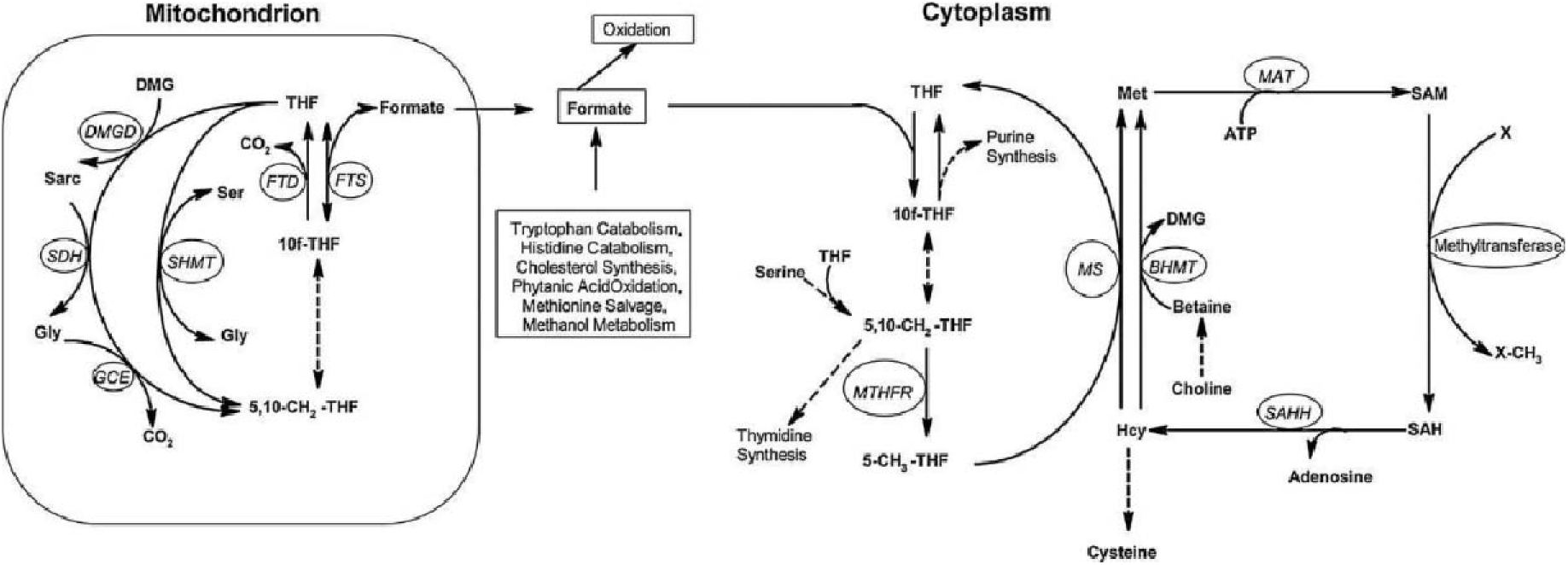
Outline of folate-mediated one-carbon metabolism [45].

To test this hypothesis, we conducted a large-scale CG experiment of Arctic × brook charr hybrids to examine how nutritional stress and dietary NT supplementation influence both growth performance and one-carbon (1C) metabolic regulation. This work follows a series of experiments conducted over the last decade to study different cyclical feeding-restriction regimes on growth metrics of salmonid species (*S. alpinus*, *S. fontinalis*) at experimental scale [22, 47] and commercial-scale [33]. The objectives of this study were to determine whether (1) a prolonged winter fasting period applied within a commercial production at the grow-out phase can enhance productivity through compensatory growth in Arctic × brook charr hybrids, (2) fasting induces measurable impairments of one-carbon (1C) metabolism consistent with vitamin B deficiency, and (3) dietary nucleotide (NT) supplementation can kick-start post-fasting growth recovery when feeding resumes by enhancing methylation capacity and sparing endogenous methyl donors. This work provides new insight into 1C metabolic adjustments during fasting and recovery in a hybrid of highly adapted charr species. It highlights the potential of NT-enriched diets within a restriction-feeding cycle as a low-risk, functional feed strategy and proposes biochemical markers that link nutrition, metabolism, and growth performance in a commercial aquaculture species.

## 2. Materials and Methods

### 1. Rearing conditions

Adult (> 350 g) all-female triploid hybrids originating from a cross between females *S. alpinus* (Fraser strain) and neomales *S. fontinalis* (Baldwin strain) were stocked and maintained at Pisciculture des Monts-de-Bellechasse inc., located in St-Damien-de-Buckland, Québec, Canada (46°37’41.5’N 70°39’08.5’W) during the months of October 2019 through March 2020 (mean temperatures: Oct, 7.2±0.4; Nov, 5.7±0.5; Dec, 5.3±0.2; Jan, 5.2±0.2; Feb, 5.1±0.3; Mar, 5.4±0.4 °C) for a total of 180 days. The natural photoperiod covered 11.4L: 12.6D to 9.5L: 14.5D. Installations were supplied with aerated freshwater from local groundwater and makeup water flow during the experiment was 100 ±10 m^3^ h^-1^, allowing two complete water turnovers per hour. Oxygen saturation levels measured daily was at least 90% and temperatures were monitored hourly using data loggers (IBWetLand, AlphaMach Inc., Sainte-Julie, QC, Canada). Fish were randomly assigned to the head sections of two concrete raceway units separated into two groups each with a meshed partition introduced directly in the raceway section within the same greenhouse building, creating four groups in duplicates (n=778 fish x 2 sections x 2 raceways). All procedures involving animals were conducted in accordance with the Canadian Council on Animal Care (CCAC) guidelines and were approved by the Université de Moncton Animal Care Committee (UdeM 19-12).

### 2. Experimental design and limitations

The experiment included two 90-day phases. During Phase 1 (experimental restriction), feeding groups were set up in duplicates and control fish groups were fed *ad libitum* with a standard diet (Orient LP 18, Skretting, St. Andrews, New-Brunswick, Canada), while restricted fish groups were not fed. During Phase 2 (exploratory recovery), all groups were fed to satiety either with the standard diet or a nucleotide-supplemented diet (Response LP 18, Skretting) spray-dried with NT from enzymatic hydrolysis of *Saccharomyces cerevisiae* fermentation extract at a dose of 1g kg_feed_^-1^. Dosage was validated with the feed company’s nutritionist. For Phase 2, replication was lost as the downstream group in both raceways received the NT-supplemented diet to assess its effect on growth and metabolism. Groups are identified as Control (Ctl), Ctl:NT, starved, and starved:NT. This was the most optimal setup agreed upon with the industry partner to accommodate the important fish biomass mobilised for diet comparison during the growth trials. Therefore, phase 2, was set as an exploratory assessment of growth metrics and data is presented without rigorous statistical analysis. In this respect, estimated mass (Mest), linearized predicted growth rate (pSGR) (See Section 2.7) and skewness coefficient of specific growth rate (SGR) (See section 3.1) calculations were conducted to provide enhanced oversight for a clearer evaluation of the growth metrics [48, 49]. The experimental design is presented in Figure 2 and includes the sampling and measurement sequence with weekly temperatures. Tissue sampling began at the end of the restriction period corresponding to day 0.

**Figure 2.**
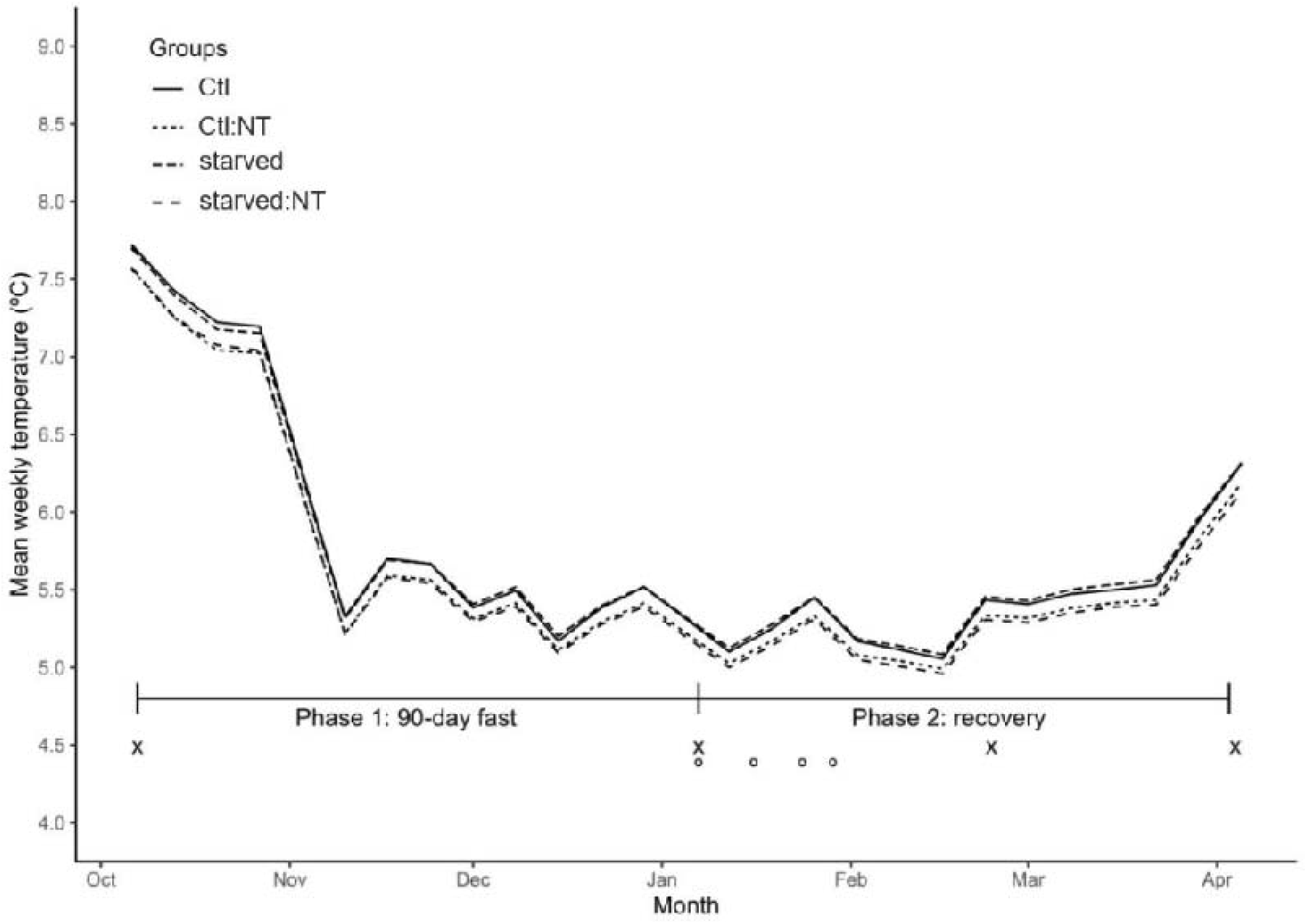
Mean weekly temperatures for all feeding regime groups, control (Ctl) and starved with or without nucleotide (NT) enriched diets throughout the trials and size measurements and tissue sampling sequence illustrated by ‘x’ and ‘o’, respectively.

### 3. Tissue and blood sampling

During phase 2, ten fish per group (n = 10) were sampled at each time point (D0, D9, D16 and D21) (n = 10 fish x 4 groups x 4 days, 160). Fish were quickly euthanized by percussive stunning, measured and weighed, and samples of blood (1.5 mL from the caudal vein) and liver were collected. Liver samples were flash-frozen on dry ice. Blood was drawn from the caudal vein into heparinized tubes, centrifuged at 2,500 × g for 10 min, and the resulting plasma transferred to clean tubes and frozen on dry ice. All samples were transported on dry ice and stored at −80 °C until analysis at the University of Moncton (Moncton, NB, Canada).

### 4. Growth measurements

Phase 1 lasted from 7 October 2019 to 7 January 2020, followed by Phase 2, which lasted until 6 April 2020. Diet groups were formed immediately after the restriction period (Phase 1). Wet body mass (g) and total fork length (mm) were recorded at 45-day intervals corresponding to the beginning, middle and end of each trial phase. On 8 October, 75 individuals per group (∼10%) were tagged intraperitoneally with passive integrated transponder (PIT) tags (12 mm 134 kHz, Biomark inc., Boise, ID, USA) for longitudinal growth analysis. All groups were fasted 24 hours prior to each measurement. On sampling days, approximately 20% of the total population for each group were transferred in batches into 1m3 tanks (Xactics Canada) at a maximum density of 150 kg m^-^3 to be lightly sedated with a 25-35 mg L-1 dose of tricaine methanesulfonate (MS-222) for 15-20 minutes prior to measurement. An automated system was used for rapid measurement and recording of wet body mass and total fork length data, as well as individual PIT tags detection (BigFin Scientific, Austin, TX, USA). The non-captured tagged fish during the sampling of 20% of the tank’s biomass, were all recovered through rapid screening of the remaining fish using rapid pit tag detection and rapidly weighed and measured to complete our individual SGR calculations based on 75 fish.

### 5. Reagents

All chemicals were of the highest purity, and HPLC grade solvents and nanopure water were used. 2,3,4,5,6-pentafluorobenzyl bromide (PFBBr) (101052-25G), hexane (34859-1L), acetone (650501-1L), sodium formate (13C, 99 %) (279412-1G), L-homocysteine (69453-50MG), S-(5’-adenosyl)-L-methionine chloride (A7007-25MG), 7-fluorobenzo-2-oxa-1, 3-diazole-4-sulfonic acid (F-4383), S-adenosylhomocysteine (A9384-25MG), and L-methionine-(methyl-D3) (98%) (300616-1G) were purchased from Sigma-Aldrich.

### 6. Metabolites analyses

Plasma formate concentrations were measured using an isotope-dilution GC-MS assay as described by Lamarre et al. [50], with slight modifications. Methionine was quantified concurrently using the same assay, referenced to a standard curve (0–400 μM) with L-methionine-(methyl-D₃) as the internal standard. The alkylation procedure was extended to 30 minutes at 60 °C to enhance methionine derivatization. Derivatized analytes were then quantified using a gas chromatograph (model 7890B; Agilent Technologies, Santa Clara, CA, USA) interfaced with a single quadrupole inert mass selective detector (MSD, model 5977B). Circulating homocysteine concentration was quantified by HPLC following Vester and Rasmussen [51] using an Agilent 1100 series HPLC (Agilent Technologies, Santa Clara, CA, USA) equipped with a fluorescence detector module (G1321A). The concentrations of S-adenosylmethionine (SAM) and S-adenosylhomocysteine (SAH) in livers were measured by HPLC as described by Jacobs et al. [52].

### 7. Calculations

#### Specific growth rate (SGR)

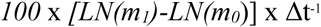

where m_1_ is the final mass of individual fish, m_0_ is the initial mass of individual fish, and Δt the time elapsed between m_1_ and m_0_.

*Predicted specific growth rate* or *pSGR* as described by Elliot and Hurley [48]

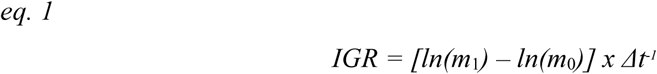

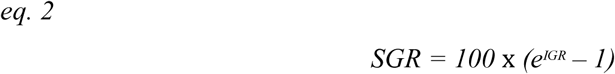

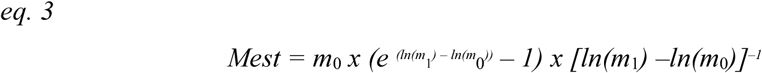

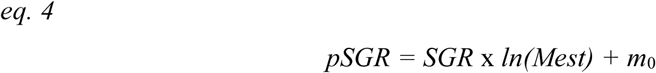

where m_1_ is the final mass of individual fish, m_0_ is the initial mass of individual fish, and Δt the time elapsed between m_1_ and m_0_; IGR is instant growth rate, Mest is estimated mass between Δt, and pSGR is linearized predicted growth rate.

#### FCE

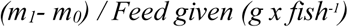

where m*_1_*is mean final mass of a group, m*_0_* is initial mean mass of a group and feed given is in dry weight (g / fish)

### 8. Statistical analyses

All statistical analyses were performed using R statistical software (version 3.5.2, CRAN, Vienna Austria). Data are expressed as means ± standard deviations (SD). Linearity assumptions were assessed with a *Levene* test of the *car* package [53]. For Phase 1, a mixed-effect ANOVA was conducted to evaluate the effects of restriction treatment (control vs. starved) on wet body mass and total fork length, with fish tank as a random factor nested within the fixed factor of treatment. Due to the lack of replicated raceway units during Phase 2, morphometric analyses are exploratory and do not support formal treatment-level inference of group means. PIT-tagged fish were used for longitudinal analysis of SGR% and the Fisher-Pearson coefficient of skewness was calculated using the *e1071* package [54] to evaluate shift trends in recruitment of fish for feeding associated with fasting and nucleotide (NT) supplementation. Due to the absence of replicate tanks in Phase 2, treatment-level statistical inference is limited. However, to explore inter-individual metabolic responses, one-carbon metabolite data was analyzed at the individual fish level. This approach is consistent with observations in metabolomics studies, where biological signal is often driven by individual variability rather than treatment means [55]. A mixed-effects ANOVA was used as an exploratory tool, treating fish as the observational unit, with post hoc comparisons performed using Tukey’s Honest Significant Difference (TukeyHSD) test at a 95% confidence level.

## 3. Results

### 3.1. Growth assessment

At the onset of the experiment, initial wet body mass and total fork length (TFL) did not differ significantly among groups (P > 0.05). By day 45, starved fish exhibited significant size reductions relative to controls, with wet body mass decreasing by 22% (0.78-fold; P = 0.004) and TFL by 3% (0.97-fold; P = 0.045). At the end of Phase 1, the size disparity had further increased, with starved fish showing a 1.43-fold lower body mass than controls (P < 0.001), corresponding to a difference of 249.5 g. Cumulatively, starved individuals lost an average of 36.6 g in body mass during the trial, representing a 5.2% reduction, whereas fed groups gained 210.7 g (53.3%). Data on fish size during Phase 1 (the 90-day restriction period) are presented in Table 1. Negative growth was consistently observed among all PIT-tagged fish in the starved group, as indicated by a calculated specific growth rate (SGR) in mass of –0.15 % per day.

**Table 1.**
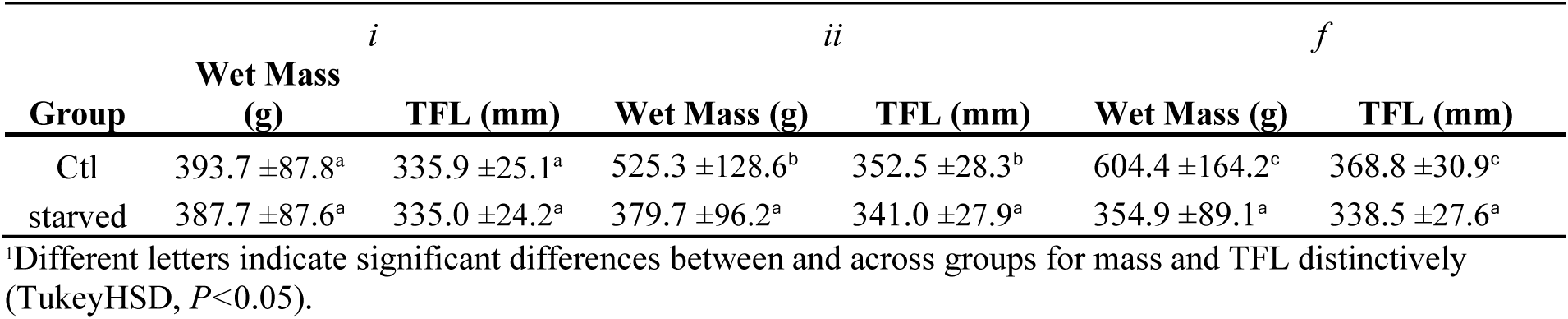
Wet body mass and total fork length (TFL) of fish during the restriction period (Phase 1) measured initially (i), 45 days later (ii) and at the end of the restriction (f) (n = 150 group-1 time-1).

During Phase 2 (the 90-day recovery period), the starved:NT group exhibited the greatest growth response, with a 1.76-fold increase in body mass. This compared to 1.67-fold in the starved group, 1.32-fold in Ctl:NT, and 1.24-fold in the Ctl group (Table 2). Nucleotide (NT) supplementation enhanced growth in both NT-fed groups and final body masses was 10.4% and 3.7% higher in starved:NT and Ctl:NT groups relative to their respective non-supplemented counterparts. Changes in TFL followed the same overall trends.

**Table 2.** Wet body mass, total fork length (TFL) measured after restriction (D0) and recovery (D90), and feed conversion efficiency (FCE), food intake (FI, g fish-1), and amount given (kg) measured at day 90 after formation of fish groups control (Ctl) and starved, refed with (:NT) or without NT for the growth assessment (n = 150 ±10 individuals group-1 time-1).

| Groups | Wet mass (g) |  | TFL (mm) |  | FCE | FI | Feed (kg) |
| --- | --- | --- | --- | --- | --- | --- | --- |
|  | D0 | D90 | D0 | D90 |  |  |  |
| Ctl | 611.3 ±177.6<sup>b</sup> | 759.4 ±209.6 | 371.1 ±32.3<sup>b</sup> | 394.5 ±35.2 | 0.521 | 284.03 | 222.9 |
| Ctl:NT | 596.3 ±146.9<sup>b</sup> | 787.0 ±207.5 | 366.0 ±29.3<sup>b</sup> | 393.2 ±32.2 | 0.597 | 319.21 | 233.7 |
| starved | 346.4 ±101.7<sup>a</sup> | 577.4 ±158.4 | 337.4 ±29.0<sup>a</sup> | 368.9 ±32.3 | 0.778 | 296.75 | 225.7 |
| starved:NT | 361.9 ±77.1<sup>a</sup> | 637.3 ±143.1 | 339.3 ±26.5<sup>a</sup> | 373.8 ±27.6 | 0.794 | 346.83 | 240.3 |

<sup>1</sup>Means followed by different letters are significantly different (TukeyHSD, $P<0.05$ ).<sup>2</sup> Statistical analyses were only performed on restriction regime replicates at D0.

Starvation led to marked improvement in feed conversion efficiency (FCE) in both starved groups, with a pooled 1.42-fold increase compared to control groups. The starved:NT group exhibited the highest FCE overall, surpassing both Ctl and Ctl:NT groups by 1.52- and 1.33-fold, respectively, and slightly outperforming the other starved group by 1.03-fold. NT supplementation also promoted feed intake: fish in the Ctl:NT and starved:NT groups consumed 35–50 g more feed per individual than their respective standard diet-fed counterparts (Table 2). Total feed distributed was nearly identical between Ctl and Ctl:NT groups (475.9 and 476.6 kg, respectively), while the starved and starved:NT groups received 47.4% and 50.5% of the control group’s feed allocation.

Specific growth rate (SGR) analysis revealed clear compensatory mass gain in both starved groups. By Day 45 of recovery, SGR values in these groups were nearly double those of Ctl groups, regardless of NT supplementation (Table 3). NT further enhanced this response, the starved:NT group showed a 1.83-fold higher SGR than Ctl at Day 45, and a 2.31-fold increase by Day 90, i.e., at the end of Phase 2. Overall, NT supplementation improved SGR in both feeding regimen groups relative to their corresponding non-supplemented controls.

**Table 3.**
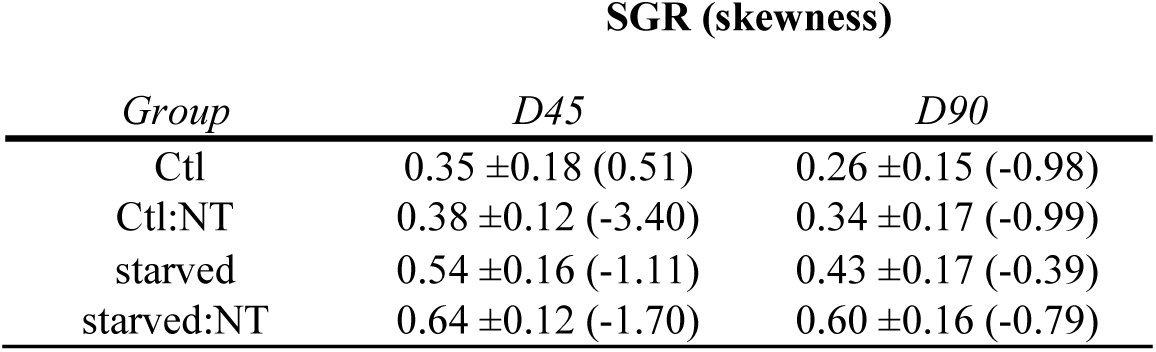
Specific growth rate (SGR) of individually tagged fish distributed in the four diet regime groups (control (Ctl) and starved groups refed with (:NT) or without NT) during refeeding with skewness data for SGR calculated with t<u>he Spearman rank correlation in parenthesis. (n = 70 fish grou</u>p-1 time-1).

Despite accelerated growth, full catch-up was not achieved within the 90-day recovery period. The starved:NT group exhibited the smallest final body-mass gap relative to the Ctl group, with a 1.16-fold difference, corresponding to 122 g at the end of recovery. Specific growth rates (SGR) observed in the starved groups at the end of Phase 2 were integrated into a linearized growth-trajectory prediction model [48]. The model predicted specific growth rates (pSGR) that were 2.1-fold and 1.7-fold higher than the Ctl group in the Starved and Starved:NT groups, respectively (Figure 3). These projections indicate that full catch-up in body mass would occur after 211 days for the Starved group and 125 days for the starved:NT group, at which points predicted mean body masses would reach 1.09 kg and 0.85 kg, respectively.

**Figure 3.**
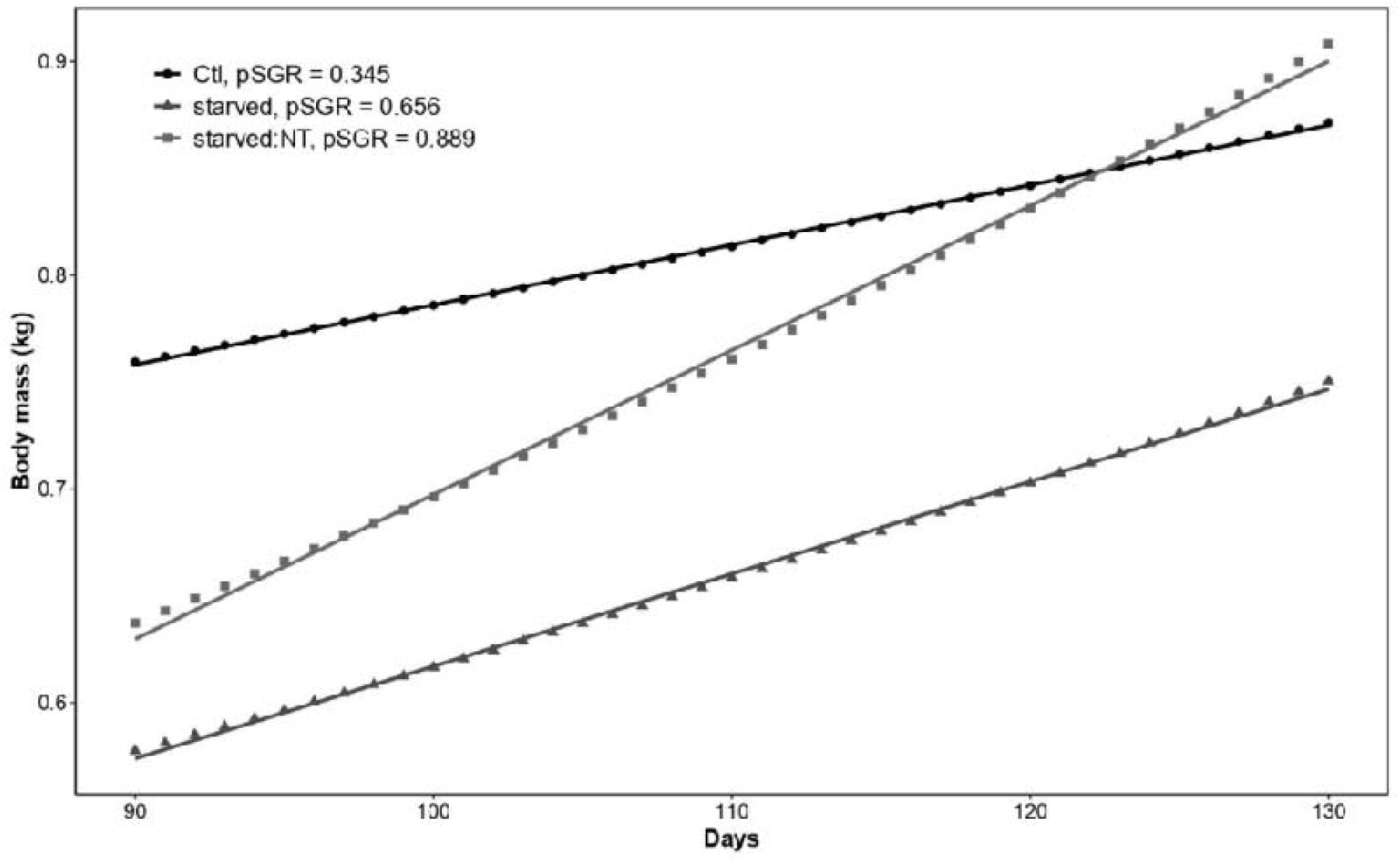
Estimated growth trajectories after day 90 of refeeding (Phase 2) for continuously fed (Ctl), previously starved refed with (starved:NT) or without nucleotide groups based on predicted specific growth rate (pSGR) calculated as described by Elliot and Hurley [48].

Skewness of SGR among tagged individuals was analyzed to evaluate intra-group variation in feed access and growth performance. After Phase 1, the Control group exhibited an approximately normal growth distribution (skewness = −1.01), whereas both starved groups showed a pronounced right-skewed distribution (skewness = 10.08), consistent with disrupted and heterogeneous growth resulting from starvation (data not shown). By day 45 of Phase 2, the growth distributions of the starved groups had shifted toward normality (skewness range: −1.70 to −0.39; Table 3), indicating a more uniform recovery. Noteworthy, NT supplementation in the Ctl:NT group initially induced a left-skewed distribution (skewness = −3.40) at day 45, which normalized by day 90. Final skewness values indicate more balanced feed intake and homogenous growth distribution among previously starved fish (average skewness = −0.59) compared to the continuously fed groups (−0.98).

### 3.2. Key metabolites of one carbon metabolism

At the end of Phase 1, continuously fed fish from the Ctl groups had significantly greater body mass (624.1 ±137.8 g) than starved fish (398.0 ±121.9 g; p=0.032). Circulating formate concentrations were significantly higher in starved fish at the end of Phase 1, showing a 1.28-fold increase compared to control (25.16±13.24 vs. 32.21±10.25 µM, p = 0.041) (Figure 4). During the recovery period, the feeding regime alone did not affect formate concentrations. However, diet had a marked effect on formate concentrations in previously starved groups: NT-fed individuals displayed significantly elevated formate concentrations at Day 9 (p<0.001), Day 16 (p=0.039) and Day 21 (p<0.001) compared with those fed the standard diet. Average formate concentrations across the recovery phase were comparable between the two NT-fed groups (Ctl:NT = 44.16 μM and starved:NT = 44.19 μM), and lower in the standard-diet groups (Ctl = 22.58 μM and starved = 27.79 μM). Overall, NT supplementation increased formate concentrations by 1.76-fold in continuously fed groups and by 1.37-fold in previously starved fish.

**Figure 4.**
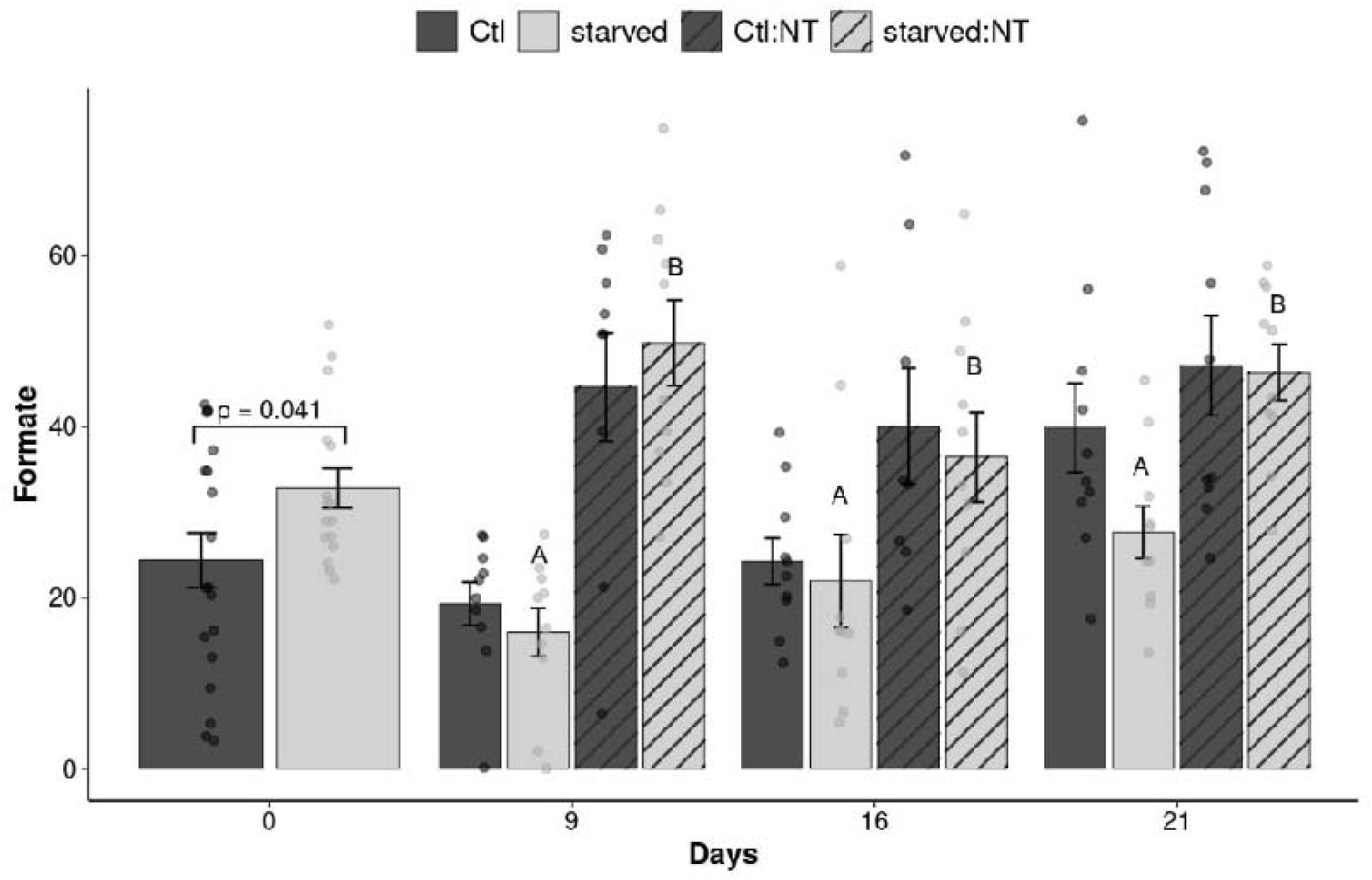
Formate concentrations (µM) measured in plasma of charr hybrid (*S. alpinus* x *S. fontinalis*) individuals of continuously fed control (Ctl), previously starved (starved) groups refed with (Ctl:NT; starved:NT) or without nucleotide (NT) enriched diets following a 90-day winter fast (Day 0) and during subsequent refeeding days. Caps and lowercase letters indicate differences between diet within feeding regime groups, and between feeding regime within diet groups, respectively. The absence of letters indicates non-significant effect. Bars indicate mean ± SD, jittered dots, individual fish data with outliers hidden for clarity.

At the end of Phase 1, S-adenosylmethionine (SAM) concentrations were significantly higher in the Ctl group than in starved fish (estimate = 865.6; p = 0.018), whereas S-adenosylhomocysteine (SAH) concentrations did not differ significantly (p = 0.161) (Table 4). Within nine days of refeeding, SAM concentrations in previously starved groups had returned to control levels and remained comparable across all treatments for the remainder of the recovery phase. On Day 9 of Phase 2, SAH concentrations were significantly lower in starved fish fed the standard diet compared with both Ctl (p = 0.0256) and Ctl:NT (p = 0.0502) groups. Analysis of the SAM:SAH ratio following Phase 1 revealed markedly higher values in continuously fed fish (Ctl = 6.56 ± 2.23) relative to starved fish (3.79 ± 1.28; p < 0.001) (Figure 5), indicating a reduced methylation potential in starved individuals.

**Figure 5.**
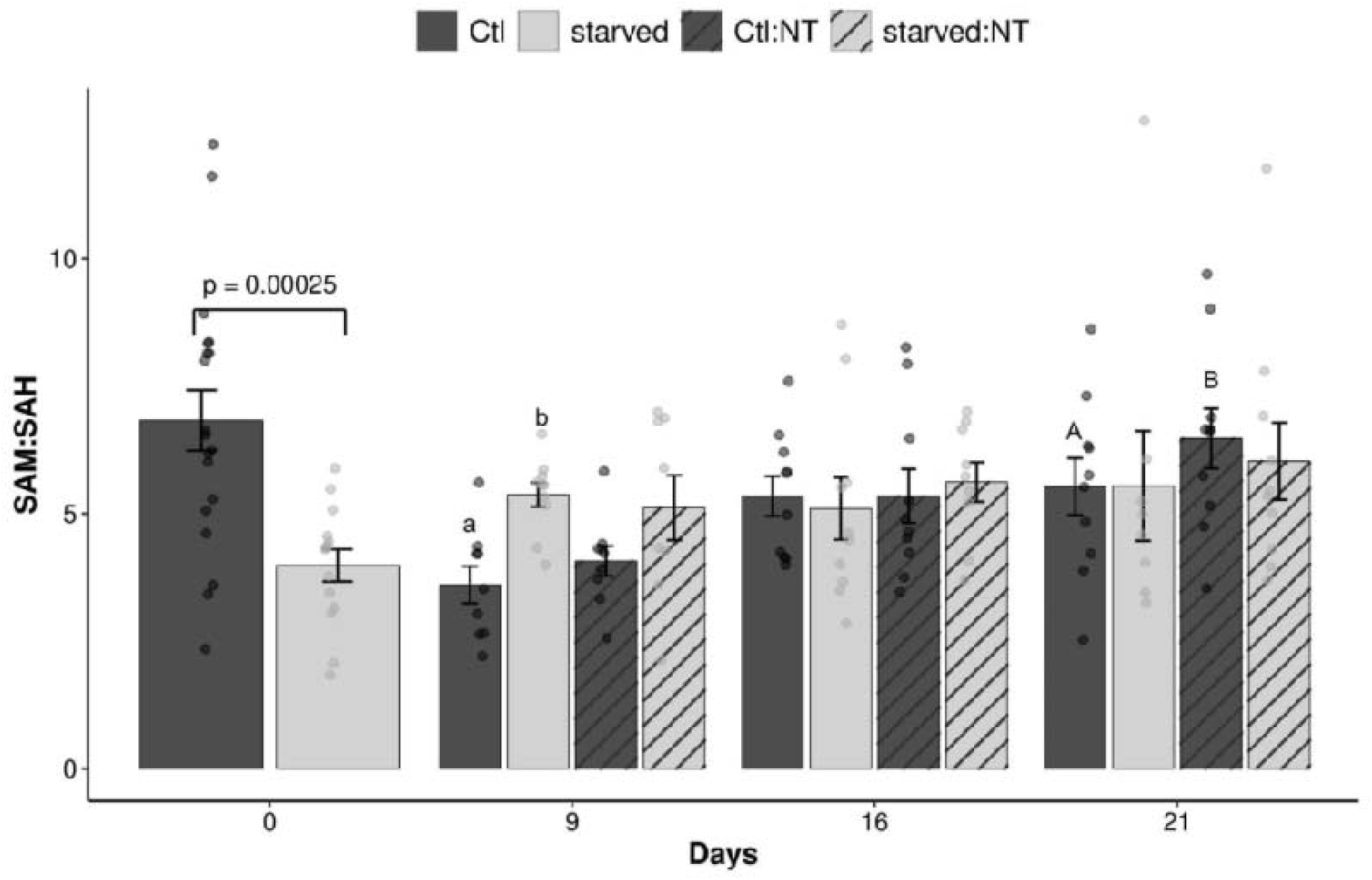
S-adenosylmethionine (SAM): S-adenosylhomocysteine (SAH) ratio measured in liver tissue of charr hybrid (*S. alpinus* x *S. fontinalis*) individuals of continuously fed control (Ctl), previously starved (starved) groups refed with (Ctl:NT; starved:NT) or without nucleotide (NT) enriched diets following a 90-day winter fast (Day 0) and during subsequent refeeding days. Caps and lowercase letters indicate differences between diet within feeding regime groups, and between feeding regime within diet groups, respectively. The absence of letters indicates non-significant effect. Bars indicate mean ± SD, jittered dots, individual fish data with outliers hidden for clarity.

**Table 4.**
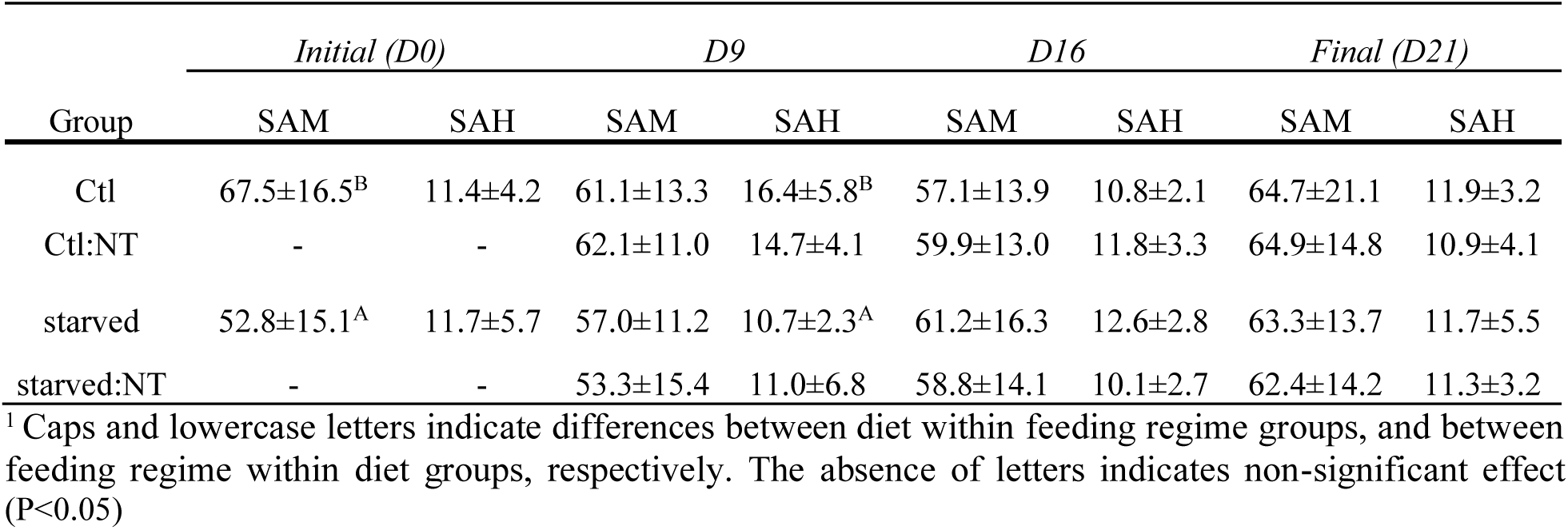
Liver S-adenosylmethionine (SAM) and S-adenosylhomocysteine (SAH) values expressed in µM g tissue^-1^ of fish sampled at the end of the restriction period (D90), 9-, 16- and 21-days post-refeeding (n = 10 group^-1^ time^-1^).

|  | *Initial (D0)* |  | *D9* |  | *D16* |  | *Final (D21)* |  |
| --- | --- | --- | --- | --- | --- | --- | --- | --- |
| Group | SAM | SAH | SAM | SAH | SAM | SAH | SAM | SAH |
| Ctl | 67.5±16.5<sup>B</sup> | 11.4±4.2 | 61.1±13.3 | 16.4±5.8<sup>B</sup> | 57.1±13.9 | 10.8±2.1 | 64.7±21.1 | 11.9±3.2 |
| Ctl:NT | - | - | 62.1±11.0 | 14.7±4.1 | 59.9±13.0 | 11.8±3.3 | 64.9±14.8 | 10.9±4.1 |
| starved | 52.8±15.1<sup>A</sup> | 11.7±5.7 | 57.0±11.2 | 10.7±2.3<sup>A</sup> | 61.2±16.3 | 12.6±2.8 | 63.3±13.7 | 11.7±5.5 |
| starved:NT | - | - | 53.3±15.4 | 11.0±6.8 | 58.8±14.1 | 10.1±2.7 | 62.4±14.2 | 11.3±3.2 |

<sup>1</sup> Caps and lowercase letters indicate differences between diet within feeding regime groups, and between feeding regime within diet groups, respectively. The absence of letters indicates non-significant effect (P<0.05)

During recovery, the feeding regime significantly influenced the SAM:SAH ratio. On day 9, previously starved fish refed with the standard diet exhibited significantly higher SAM:SAH ratios compared to continuously fed Ctl fish (p=0.016), suggesting a transient compensatory adjustment in methylation capacity. A significant dietary effect was also observed at day 21 in both continuously fed groups: fish in the Ctl:NT group displayed higher SAM:SAH ratios (p=0.040) than those in the Ctl group.

At the end of Phase 1, homocysteine (Hcy) concentrations were significantly elevated in starved fish compared to Ctl (1.21 ±1.10 µM vs 0.30 ±0.43µM, p< 0.001, Figure 6), indicating a disruption in one-carbon metabolism. However, during the recovery period, neither diet nor feeding regime significantly affected Hcy concentration in either control or starved groups. Mean Hcy values remained similar between feeding treatments (Ctl: 0.41 ± 0.55 µM; starved: 0.38 ± 0.71 µM), and between diet groups (NT: 0.48 ±0.69 µM; Std: 0.25 ± 0.51 µM), with no significant time effect (p = 0.611). In many samples, Hcy were near or below the detection limit, and high variability likely limited statistical power.

**Figure 6.**
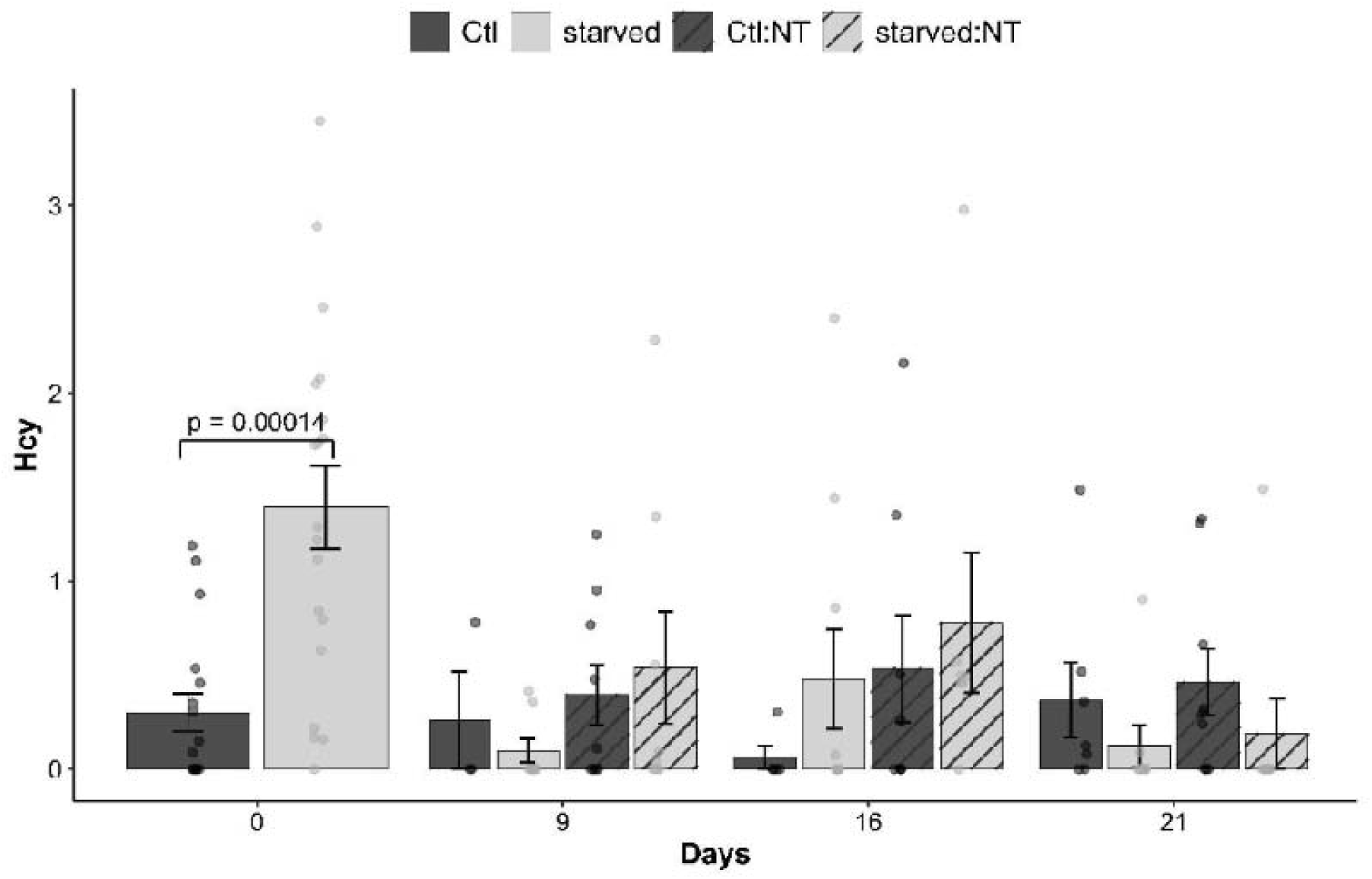
Homocysteine (Hcy) concentrations (µM) measured in plasma of charr hybrid (*S. alpinus* x *S. fontinalis*) individuals of continuously fed control (Ctl), previously starved (starved) groups refed with (Ctl:NT; starved:NT) or without nucleotide (NT) enriched diets following a 90-day winter fast (Day 0) and during subsequent refeeding days. Caps and lowercase letters indicate differences between diet within feeding regime groups, and between feeding regime within diet groups, respectively. The absence of letters indicates non-significant effect. Bars indicate mean ± SD, jittered dots, individual fish data with outliers hidden for clarity.

Methionine (Met) concentrations were significantly reduced by starvation, with lower values in starved fish compared with controls at the end of Phase 1 (52.77 ± 5.85 µM vs. 84.90 ± 41.18 µM; p = 0.0024; Figure 7). During the recovery phase, the feeding regime alone had no significant effect on Met concentrations. However, NT supplementation significantly increased circulating Met levels, with higher concentrations observed in the starved:NT group on Day 9 (p = 0.0263) and in both starved:NT (p = 0.0314) and Ctl:NT (p = 0.0237) groups on Day 21.

**Figure 7.**
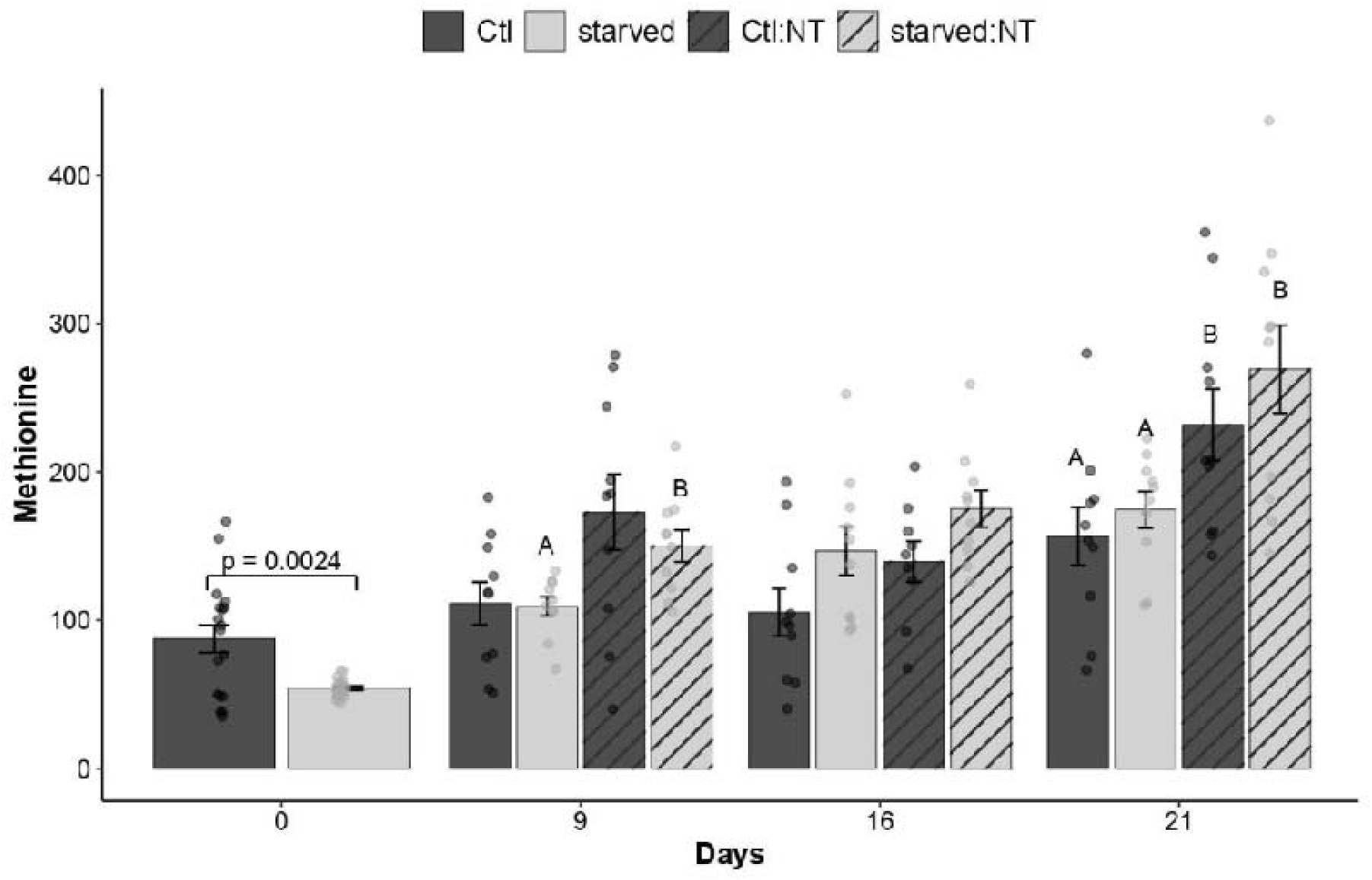
Methionine (Met) concentrations (µM) measured in plasma of charr hybrid (*S. alpinus* x *S. fontinalis*) individuals of continuously fed control (Ctl), previously starved (starved) groups refed with (Ctl:NT; starved:NT) or without nucleotide (NT) enriched diets following a 90-day winter fast (Day 0) and during subsequent refeeding days. Caps and lowercase letters indicate differences between diet within feeding regime groups, and between feeding regime within diet groups, respectively. The absence of letters indicates non-significant effect. Bars indicate mean ± SD, jittered dots, individual fish data with outliers hidden for clarity.

## 4. Discussion

This study examined how a protracted winter starvation and a spring refeeding sequence influenced growth performance and one-carbon (1C) metabolism in Arctic charr × brook charr hybrids, and whether dietary nucleotide (NT) supplementation can promote methyl-unit sparing to support compensatory growth. Consistent with the exceptional starvation tolerance of *Salvelinus* species, the fish endured a 90-day feed deprivation period with no mortality, marginal body mass reduction and recovered rapidly upon resumption of feeding. Starvation induced moderate changes in metabolic and methylation indices, including elevated circulating formate and homocysteine concentrations and reduced methionine and SAM:SAH ratios, indicating a temporary attenuation of 1C-mediated methylation capacity. Refeeding restored these parameters within weeks, demonstrating a strong capacity for metabolic re-equilibration.

Data from this study highlight a consistent stimulatory effect of dietary nucleotides on mitochondrial one-carbon flux. NT supplementation further enhanced recovery, increasing methionine availability, formate concentrations, and overall methylation potential in both feeding groups. These responses are consistent with improved 1C flux and anabolic readiness [56]. While the present results do not demonstrate a direct mechanistic link, they suggest a functional association between 1C metabolic status and growth recovery following fasting. This potential relationship warrants further investigation to determine whether dietary nucleotides effectively spare methyl donors or modulate 1C flux at the enzymatic level. Collectively, these findings point to a promising nutritional approach to optimize refeeding efficiency and promote sustainable production in charr aquaculture.

### 4.1 Growth assessment

Building on the work of Imsland and Gunnarsson [57], and more recently Le François et al. [22] and Drouin-Johnson et al. [33], our study provides a first assessment of the growth response of Arctic charr × brook charr hybrids following a winter restriction-spring refeeding cycle. The hybrid displayed good tolerance to fasting, with only a 5.2 % reduction in body mass after a 13-week period. The negative growth observed confirmed the successful induction of a prolonged catabolic condition i.e. lipid mobilization Phase II proposed by Bar and Volkoff [6], which is required for a full expression of a compensatory growth response. During refeeding, starved fish exhibited specific growth rates nearly twice those of continuously fed groups, and NT supplementation further enhanced both growth and feed intake. Although full catch-up was not achieved within the 90-day recovery period, a clear compensatory trajectory unfolded. Based on growth projections, full compensation was estimated at 125 days, which is comparable to previous estimates for Arctic charr: 117 days in Imsland and Gunnarsson [23] and 86 days in Le François et al. [22]. The slightly longer catch-up time calculated for the hybrid may be the consequence of its larger body size and older age, factors known to reduce the plasticity of metabolic reorganization [62].

In terms of loss of body mass during feed restriction, Atlantic salmon (*Salmo salar*) generally loses body mass more rapidly with reported reductions of 5-11.3% over periods ranging from 6 to 12 weeks [58–61]; brook charr juveniles (80-120 g initial mass) lost 18% body mass after a 28-day winter fast [47] and Arctic charr juveniles (Nauyuk strain, 150-200 g) maintained body mass over a 14.5 week fast [22]. These contrasting responses underscore the exceptional fasting tolerance of the Arctic charr and our hybrid. However, due to the lack of experimental replication during the recovery phase, results should be interpreted with caution.

### 4.2 Growth variability and aquaculture implications

Growth variability has long been identified as a major obstacle to the productivity of charr aquaculture, leading to suboptimal feeding, increased aggression, and frequent, stressful, and costly grading operations [63, 64]. Strategies that reduce size variability within-cohort are therefore of strong interest to producers. Recent studies have shown that feed restriction reduces growth disparity upon refeeding, with more severe restrictions leading to greater uniformity [22, 65]. Our results support this trend, demonstrating that a winter fast followed by refeeding can enhance growth homogeneity in charr hybrids. To assess intra-group variability, we applied the skewness coefficient of SGR, as proposed by Hatlen et al. [49], who demonstrated that this metric can better capture disproportionate feed intake following severe feed restriction than the traditional calculation of the coefficient of variation. In our study, skewness shifted from highly positive values following starvation to near-zero values after 45 days of refeeding, indicating a rapid normalization of growth distribution. Data on skewness of SGR in tagged *S. alpinus* juveniles (180 g) retrieved from Le François et al. [22] indicate a similar pattern where control fish displayed normal growth (skewness = 1.13) while fish starved for 102 days exhibited a right-skewed distribution of 5.76, which by day 56 into refeeding had shifted toward normality (skewness = 0.137). These findings suggest that even relatively short fasting periods can help equalize subsequent growth trajectories and reduce competitive disparities among fish cohorts and the frequency of grading procedures common in charr cultivation.

Beyond growth uniformity, reduced feed input during the restriction phase represents a key opportunity for improving sustainable profitability in aquaculture. Decreased feed use not only lowers costs but also reduces the environmental footprint associated with nutrient discharge. In intensive systems, 60–80% of dietary phosphorus is released into the environment [66], contributing to eutrophication of the receiving freshwater ecosystems. Although direct measurement of phosphorus discharge was not possible in this trial, largely due to shared discharge points and flow-through dilution, feed restriction is expected to reduce total phosphorus loading proportionally to reduced feed intake. From a practical standpoint, feed restriction cycling—particularly when paired with functional additives like nucleotides upon refeeding—offers a potentially promising strategy for improving production efficiency while supporting environmental sustainability. Predictive models based on feed composition and species-specific nutrient retention rates could serve as useful tools to estimate phosphorus outputs under varying feed regimes [67]. As nutrient discharge regulations increasingly influence site licensing and expansion in countries like Canada, such data could strengthen the case for integrating feed management into sustainable intensification strategies.

### 4.3 Effect of starvation on one-carbon metabolism

One-carbon (1C) metabolism supports essential cellular functions, including nucleotide synthesis and amino acid interconversion [37]. These pathways are particularly relevant to growth and development, as demonstrated in mammalian studies. For instance, Cook et al. [68] reported severely reduced growth in rats fed a methyl-deficient diet lacking choline and substituting methionine with homocysteine (Hcy). While the direct effects of 1C deficiency on protein synthesis remain unclear in fish, associated growth impairments have been well documented [68, 69]. Beyond biosynthesis, 1C metabolism is increasingly recognized for its role in DNA methylation and epigenetic regulation, processes that influence lipid metabolism [70], phenotypic plasticity, and disease susceptibility [43]. As noted by Le François et al. [22] and Drouin-Johnson et al. [33], feed-restriction models should incorporate assessments of adiposity, since lipid reserves define the timing of metabolic and catabolic transitions during prolonged fasting [6]. In Arctic charr, known for their high lipid storage capacity, extended fasting may therefore delay shifts into strongly catabolic states. This interpretation aligns with our findings: despite 13 weeks of feed deprivation, charr hybrids exhibited only mild perturbations in 1C metabolism. This must be interpreted with caution as no normal values of the measured metabolites in salmonids were available at the time of publication. Plasma homocysteine increased approximately fourfold, a change similar in relative magnitude but much lower in absolute concentration than that observed in folate-deficient juvenile rats [45]. Formate concentrations were modestly elevated (1.28-fold), and methionine concentrations decreased significantly but not dramatically. These metabolites levels remained within the ranges reported for rats [46] and humans [50], suggesting moderate metabolic stress rather than severe disruption. Hepatic SAM:SAH ratio decreased by ∼40 % in starved fish, consistent with reduced methylation potential and decreased availability of methyl donors for DNA and protein methylation. Taken together, these biochemical patterns indicate a modest impairment of 1C metabolism during fasting, but the physiological impact appears limited—likely mitigated by metabolic suppression and efficient energy conservation in charr hybrids. Overall, these results provide novel insight into how energy-saving strategies in fish interact with cellular biosynthetic capacity under starvation in highly adapted species.

### 4.4 Effects of nucleotide supplementation on one-carbon metabolism

Upon refeeding, NT supplementation increased circulating formate concentrations in both previously starved and continuously fed fish. Formate plays a key role in one-carbon metabolism as a primary donor of methyl groups for the biosynthesis of purine and pyrimidine [34, 71]. The observed elevation in plasma formate may therefore reflect a conservation of endogenous one-carbon units facilitated by the exogenous supply of nucleotides.

This interpretation is further supported by the concurrent increased circulating methionine concentration and hepatic SAM:SAH ratio in NT-supplemented groups by day 21 of refeeding. A higher SAM:SAH ratio indicates enhanced methylation capacity, suggesting that nucleotide supplementation helped restore one-carbon metabolic function more rapidly than refeeding alone. While the effect on SAM concentration was somewhat variable, SAH levels remained stable across treatments, reinforcing the idea that NTs supported methyl donor availability without substantially altering turnover. These biochemical adjustments may underlie the improved growth performance observed in NT-supplemented fish. Enhanced 1C flux has been shown to support protein synthesis and cell proliferation in other fish models [72], which aligns with the greater growth rates recorded at Day 45. We propose that NT supplementation during the early refeeding period may have initiated a transient metabolic priming, facilitating accelerated somatic growth as recovery progressed. Because early-stage responses were expected to dominate the metabolic signal, measurements of 1C metabolites were limited to the first 21 days. Although this precludes direct correlation with later growth metrics, the timing of biochemical changes supports a plausible association between methylation recovery and compensatory growth. Future studies should extend temporal sampling and integrate transcriptomic and isotopic approaches to better resolve the dynamics and metabolic fates of one-carbon units during compensatory growth. From an applied standpoint, dietary NT supplementation appears to enhance methylation capacity and may support post-fasting tissue regeneration and growth. These findings underscore the potential of functional feeds to improve production recovery and metabolic resilience in aquaculture.

## 5. Conclusion

This study evaluated the effects of a 90-day winter fasting period followed by refeeding in Arctic charr × brook charr hybrids, with a focus on growth performance and one-carbon (1C) metabolic adjustments. The mild changes observed in circulating homocysteine, methionine, and hepatic SAM:SAH ratio suggest a moderate impairment of 1C metabolism during prolonged starvation, with limited impact on methylation capacity. Refeeding rapidly restored metabolic balance, and dietary nucleotide (NT) supplementation appeared to enhance early metabolic recovery, potentially priming fish for accelerated growth later in the refeeding period. While our findings support a role for methyl group sparing in the observed growth benefits, the absence of metabolite measurements during the peak of compensatory growth limits definitive conclusions. Given the central role of sulfur-containing amino acids (e.g., methionine, cysteine) in 1C metabolism, future studies should also examine how vitamin B status influences the transsulfuration pathway and downstream effects on growth. These results provide new insights into the metabolic flexibility of charr hybrids and highlight the potential of NT-enriched diets to support recovery and performance in aquaculture settings.

## Credit author statement

**Charles Drouin-Johnson**: Methodology, Validation, Formal Analysis, Investigation, Data Curation, Writing-Original Draft. **Nathalie R. Le François**: Conceptualization, Methodology, Visualization Validation, Investigation, Resources, Writing-Reviewing & Editing, Supervision, Project administration, Funding acquisition. **Grant W. Vandenberg**: Supervision, Funding acquisition. **Moïse Cantin**: Investigation, Resources. **Simon G. Lamarre**: Conceptualization, Methodology, Validation, Investigation, Resources, Supervision, Writing-Reviewing & Editing.

## Declaration of Competing Interest

The authors declare no competing interests.

## Funding

This research was funded by the Innovamer program of the Ministère de l’agriculture, des pêches et de l’alimentation du Québec (MAPAQ) awarded to NRLF and Mitacs (Project no. IT 15450 to NRLF, GWV). Ressources Aquatiques Québec (RAQ) provided the financial support for inter-laboratory exchange expenses of MSc candidate CDJ.

## Data Availability

Data generated or analyzed during this study will be available from the dataverse repository Borealis upon publication using this link: https://doi.org/10.5683/SP4/JT3OFI

## Acknowledgements

The authors want to express their gratitude to the technical staff at the Pisciculture-des-Monts-de-Bellechasse for their assistance during the conduct of the growth trials and sampling operations at their facilities.

## Notes

### Competing Interest Statement

The authors have declared no competing interest.

